# An open CAR-T single-cell atlas to enable in-depth characterization and rational engineering of CAR-T products

**DOI:** 10.1101/2025.10.11.681788

**Authors:** Sergio Camara-Peña, Paula Rodriguez-Marquez, Nuria Planell, Maria E. Calleja-Cervantes, Lorea Jordana-Urriza, Giacomo Cinnirella, Shlomit Reich-Zeliger, Paula Rodriguez-Otero, Esteban Tamariz, Idoia Ochoa, Nir Yosef, Juan R. Rodriguez-Madoz, Felipe Prosper, Mikel Hernaez

## Abstract

We built a CAR-T cell functional atlas from over one million cells across 13 studies, integrating data from patients and healthy donors. The atlas captures 11 phenotypes, links infusion product composition with clinical response, and reveals sex- and age-dependent effects, metabolic signatures, and rare ICANS-associated cells. This open-access resource provides a foundation to understand CAR-T cell function and guide the design of next-generation therapies.

## MAIN

Adoptive cell therapy strategies utilizing modified T lymphocytes with chimeric receptors (CAR-T cells) have emerged as promising treatments for several hematological diseases^1^, including acute lymphoblastic leukemia (ALL)^2,3^, B-cell non-Hodgkin lymphomas (NHL)^4,5^, and multiple myeloma (MM)^6,7^. Despite groundbreaking initial results, CAR-T therapy still faces major challenges, such as lack of long-term efficacy, tumor antigen heterogeneity, and treatment-related toxicities. Advances in single-cell RNA sequencing (scRNA-seq) have led to the generation of large datasets characterizing the molecular mechanisms that govern CAR-T cell function^8–10^. However, their isolated nature is hampering integrative cross-study analyses that would provide both holistic and deeper insights into CAR-T cell responses.

In this work we built a CAR-T cell functional atlas comprising more than 180 scRNA-seq samples, which allowed us to deepen into known CAR-T features, such as the link between memory-enriched infusion products (IP) and complete response^11,12^, or previously unnoticed differences in response across age and sex. Importantly, we were able to unravel and characterize cells linked to immune effector cell-associated neurotoxicity syndrome (ICANS) in post-infusion samples. Finally, we provide a unique set of open computational tools, such as a trained CAR-T cell atlas model within the scVI toolkit, and an intuitive web application.

To build such atlas, we compiled scRNA-seq data from 13 public studies, comprising ~1 million cells from 14 healthy donors and 102 patients with hematological malignancies (Fig. 1a), including comprehensive metadata, such as patient demographics, therapy-induced toxicities, and clinical responses (Fig. 1b, Extended Data Fig. 1 and Supplementary Tables 1,2). After rigorous quality control (Methods), 414,000 high-quality CD3^+^CAR^+^ cells were retained, 69.8% of them derived from IP (51.1% from patients, 18.7% from healthy donors), while 30.1% were post-infusion cells. Regarding response, 46.7% of the cells originated from complete responders (CR), 19.8% from non-responders (NR), and 4.9% from partial responder (PR) patients. Of note, 90.1% corresponded to CAR-T cells targeting CD19, while less than 10% targeted BCMA (4.9%) or other antigens (Fig. 1b).

**Fig. 1.**
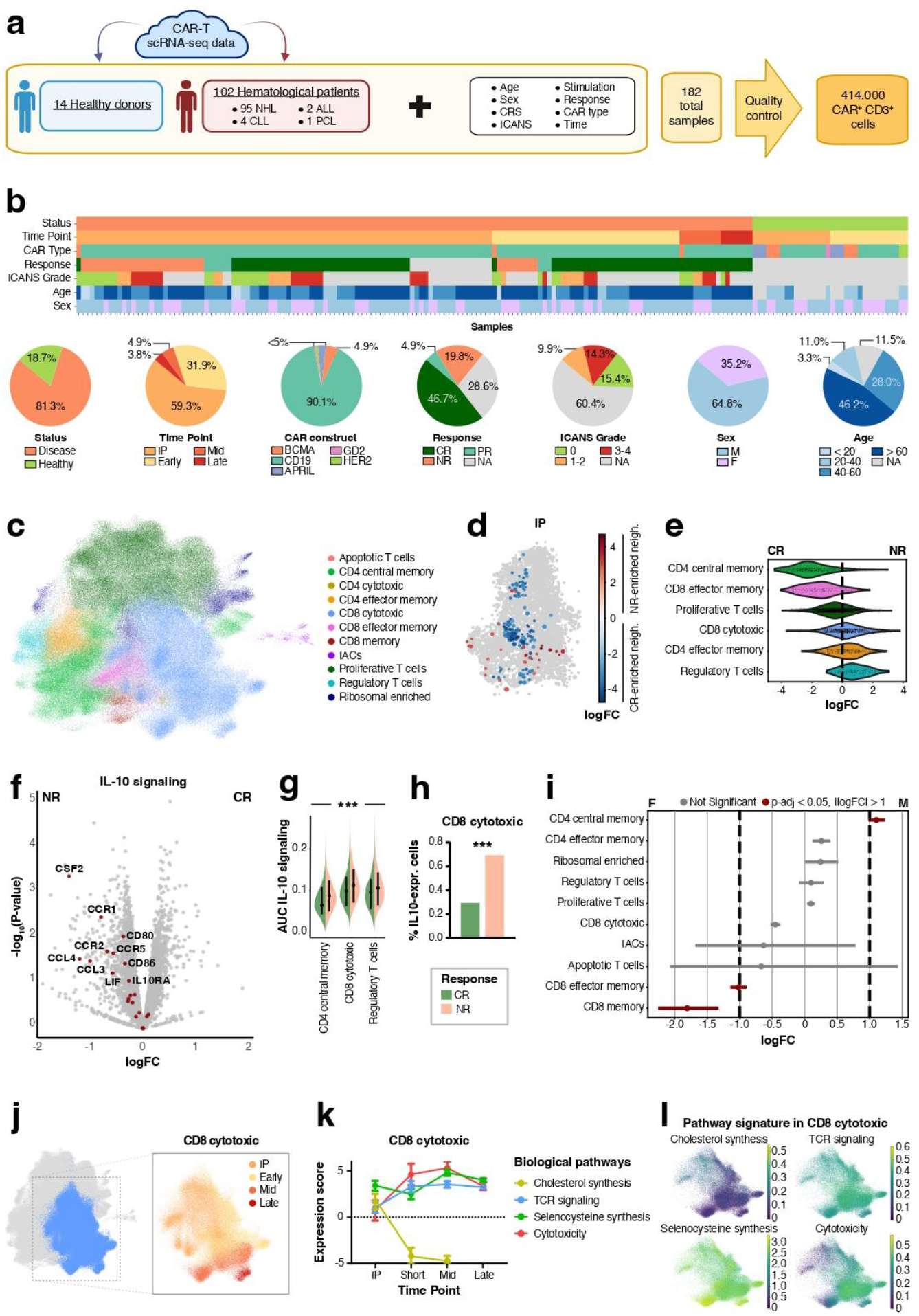
Construction and characterization of the CAR-T cell atlas. **a**. Publicly available scRNA-seq data and associated metadata from 14 healthy donors and 102 patients with hematological malignancies were integrated, yielding 182 samples encompassing 414,000 CAR^+^CD3^+^ T cells after quality control. **b**. Distribution of key metadata features across samples, including disease status, time point, CAR construct, clinical response, ICANS grade, sex, and age. Time points are categorized as infusion product (IP), early (<2 weeks), mid (2 weeks–3 months), and late (>3 months). Sex is indicated as male (M) or female (F). **c**. Complete manually annotated CAR-T cell atlas. **d**. Differential abundance analysis of IP highlighting CR-or NR-enriched neighborhoods. **e**. Neighborhood differential abundance of IP cells in CR vs. NR patients per cluster. **f**. Volcano plot showing DEGs between CR and NR/PR patients in IP. Red dots indicate IL-10 pathway genes with key genes labeled. **g**. IL-10 signaling score is increased in NR vs CR patients in CD4^+^ T_CM_, CD8^+^ cytotoxic and T_regs_. **h**. Proportion of IL-10-expressing CD8^+^ cytotoxic cells in CR and NR patients. **i**. Sex-stratified proportion analysis in CR patients. Male (M), female (F). **j**. CD8^+^ cytotoxic UMAP across time points. **k.l**. Temporal dynamics and UMAPs of pathway signatures in CR CD8^+^ cytotoxic cells. Wilcoxon test (g) and Fisher’s exact test (h) were used. ***P < 0.001.

Due to the importance of dataset integration in atlas-based analyses and following recent concerns about comparative studies and current recommendations^13^, we further evaluated six integration methods on our data. The scVI embedding was selected as it minimized batch effects while preserving biological variability (Supplementary Note 1, Extended Data Fig. 2b, Supplementary Table 3). After integration, we clustered and manually annotated 11 distinct CAR-T cell phenotypes using well-characterized genes and cell cycle markers (Extended Data Fig. 3a-c, Methods). Note that all studies contained almost all annotated T cell phenotypes within their CAR-T cells (Extended Data Fig. 3d). Overall, after this process, we obtained a highly curated, high-resolution CAR-T cell atlas (Fig. 1c).

The phenotypic composition of IPs and its relationship with clinical response is well known^11,12^. Indeed, differential abundance testing, with age and sex considered as covariates (Fig. 1d and Extended Data Fig. 4a-b, Methods), showed that IPs from CR patients were significantly enriched in memory-phenotype cells, particularly CD4^+^ central memory (T^CM^) and CD8^+^ effector memory (TEM) cells (Fig. 1e). Additionally, IPs enriched in regulatory T cells (T_regs_) were predominantly associated with NR patients (Fig. 1e). When delving into NR-enriched biological processes, we found that IL-10 signaling was upregulated across the different cell compartments, especially within CD4^+^ T_CM_, T_regs_ and CD8^+^ cytotoxic cells (Fig. 1f,g, Extended Data Fig. 4c, Supplementary Table 4,5). IL-10, primarily produced by T_regs_14, can also be expressed by CD8^+^ cytotoxic T cells during the peak of infection^10,15^. Indeed, among CD8^+^ CAR-T cells, those expressing IL-10 were significantly more abundant (OR= 2.383, p= 4.51×10^−5^) in NR patients (Fig. 1h, Supplementary Table 6), suggesting a negative influence on CAR-T functionality.

Next, we investigated the impact of age and sex in the IP composition and associated therapeutic response. Although no major sex-specific differences have been previously observed in IP composition^16^ (Extended Data Fig. 4d), a sex-stratified analysis revealed that CR-associated populations in IPs were differently enriched between men and women, with CD4^+^ T_CM_ more abundant in men and CD8^+^ TEM in women (Fig. 1i). Further stratification by age showed that the association of these populations with CR was more prominent in elder patients (>60 years) (Extended Data Fig. 4e). Interestingly, while elevated T_regs_ level generally correlate with poorer outcomes^17^, this association did not hold for elderly women (Extended Data Fig. 4e), possibly driven by hormonal^18^ or menopausal changes^19^, indicating a need for further studies. In summary, although memory-enriched IPs have been associated with CR, here we showed that distribution of memory populations is different between men and woman, being only relevant in elderly patients.

We then delved into the molecular changes that CAR-T cells undergo after infusion in patients with CR. We focused on CD8^+^ cytotoxic T cells since they accounted for 73% of post-infusion cells (note that they corresponded to 26% of IP). Thus, we analyzed the transcriptional changes undergone by this population from IP against each time point: early (<2 weeks), mid (2 weeks - 3 months) and late (>3 months) (Fig. 1j and Extended Data Fig. 5a-c; Methods). We found distinct temporal dynamics across several pathways (Fig. 1k and Extended Data Fig. 6a-f). As expected, TCR signaling and cytotoxic pathways were upregulated early after infusion (Fig. 1k,l)^20,21^. More strikingly, a sharp downregulation of cholesterol biosynthesis genes (e.g. *DHCR7, HMGCS1*, and *MSMO1*) was observed at post-infusion stages (Fig. 1k,l and Extended Data Fig. 5d). Cholesterol is known to support lipid raft organization in T cells facilitating TCR and CD3 clustering and downstream signaling^22^, while excessive cholesterol levels have been described to promote exhaustion^23^. Further studies should clarify if this reduction reflects a normalization of elevated cholesterol in IP after infusion or protective anti-exhaustion mechanisms. Interestingly, we also observed an increase in selenocysteine synthesis at mid-time (Fig. 1k,l), which supports CD8^+^ T cell proliferation and cytotoxic function while maintaining reactive oxygen species homeostasis. Note that, selenium deficiency impairs the development of functional mature T cells^24^.

To understand the molecular mechanisms associated with clinical response after CAR-T cell infusion, we next focused on CD8^+^ cytotoxic cells across patients with distinct therapeutic responses (CR, PR, and NR) (Fig. 2a,b). Among several differentially expressed processes (Extended Data Fig. 7a), key metabolic pathways were associated with response. For example, oxidative phosphorylation was downregulated in patients not achieving a CR, aligning with its role in T cell activation and persistence^25^, Wnt signaling was significantly upregulated in NR patients, consistent with its role in immune exclusion and resistance to immunotherapies^26^, and TGF-β signaling, a known immunosuppressive cytokine, was also significantly upregulated in these patients, potentially contributing to the reduction of cytotoxicity and promoting cellular exhaustion^27,28^ (Fig. 2c and Supplementary Table 7,8).

**Fig. 2.**
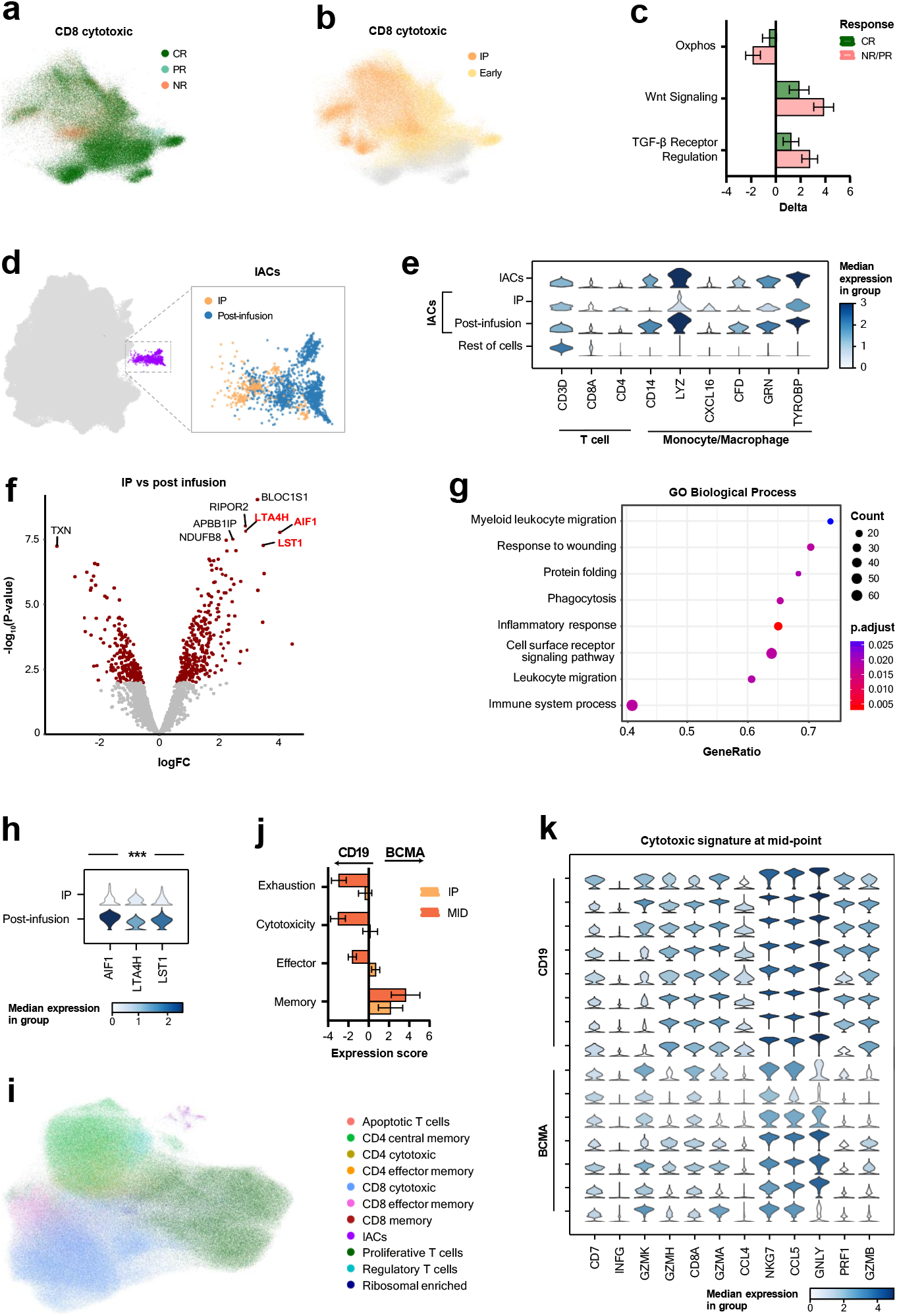
Integrative atlas-based identification of CAR-T cell molecular drivers. **a**. CD8^+^ cytotoxic cell UMAP colored by response and **b**. early time point. **c**. Oxidative phosphorylation (OXPHOS), Wnt signaling and TGF-β receptor GSEA deltas between IP and early time point in CD8^+^ cytotoxic cells from CR and NR/PR patients. **d**. ICANS-associated cells (IACs) formed a distinct cluster. Both IP and post-infusion samples are shown. **e**. Expression of T cell and monocyte/macrophage genes in IACs (IP, post-infusion) and other cells. **f**. Volcano plot showing DEGs in IACs across stages. Key genes were labeled. **g**. GSEA comparing IACs between IP and post-infusion. **h**. Expression of key DEGs in IACs between IP and post-infusion. **i**. Final UMAP after integration of new BCMA dataset into the atlas. **j**. Exhaustion, cytotoxicity, effector and memory pathway expression scores in CD19 vs BCMA CAR-T cells. **k**. Cytotoxic signature expression in CD19 vs BCMA CAR-T cells at mid time point. Mann-Whitney U test used for h. ***P < 0.001.

We identified a rare cell population co-expressing T cell (*CD3D, CD8A, CD4*) and monocyte (*CD14, LYZ*) markers (Fig. 2d,e, Extended Data Fig. 3b and 7b). While part of these cells have been previously described in their corresponding studies as ICANS-associated CAR-T cells (IACs) in the IPs^8,29^, other studies discarded this cell population probably due to their low number. Interestingly, most of the discarded cells came from post-infusion samples. In total, we identified 1495 IACs (264 in IPs and 1,231 post-infusion) (Fig. 2d and Extended Data Fig. 7c). These IAC cells exhibited upregulation of inflammatory genes (*CXCL16, CFD, GRN, TYROBP)*, with some genes *(CXCL10, CCL19*) also implicated in microglial responses to nerve injury^30–32^ (Fig. 2e and Extended Data Fig. 7d). Indeed, an upregulation of IFN-γ signaling and microglia-related pathways indicates a pro-inflammatory profile (Extended Data Fig. 7e). When comparing pre- and post-infusion states, we observed significant upregulation of inflammatory processes (*AIF1, LTA4H* genes), and IFN-γ-related genes, such as *LST1*, which reduces lymphocyte proliferation (Fig. 2f-h, Extended Data Fig. 7f and Supplementary Table 9). Additionally, after infusion, there was an upregulation of migration and myeloid functions, indicating a more pronounced monocyte-like phenotype of IAC cells after antigen recognition (Fig. 2g and Extended Data Fig. 7f). Finally, we assessed their clinical relevance and observed a statistically significant association between ICANS severity and the presence of IAC in the peripheral blood of patients increasing to >80% of the cases (OR=6.067, p=0.0497) (Supplementary Table 10).

The central goal for building a CAR-T cell reference atlas is to facilitate, integrative analyses by enabling the continuous integration of new datasets. To showcase such capabilities, we incorporated an in-house dataset comprising three additional patients treated with BCMA-targeted CAR-T therapy (Extended Data Fig. 8a and Supplementary Table 11; Methods). Our automatic integration and annotation approach leveraging the scArches-scANVI tools achieved an annotation accuracy of 82.3%, in concordance with current state-of-the-art automatic annotators (Fig. 2i). 40,848 new cells were integrated into the atlas (Extended Data Fig. 8b, allowing us to investigate differences between CAR-T cells engineered with BCMA-versus CD19-targeting constructs (Extended Data Fig. 8a-c). At mid-time after infusion, BCMA CAR-T cells showed significantly reduced expression of genes associated with effector function, cytotoxicity, and exhaustion^33^ compared to CD19 CAR-T cells (Fig. 2j,k and Supplementary Table 12). Moreover, gene ontology enrichment analysis showed that main differences were mostly restricted to general pathways, being macromolecule biosynthesis and immune response one of the most significantly affected biological processes (Extended Data Fig. 8d). However, further evaluation would be required to determine the functional relevance of these transcriptional differences as some inter-patient variability was observed (Fig. 2k). The integration of this BCMA CAR-T enriched new dataset allowed us to assess molecular differences, highlighting the applicability of the atlas for cross-study comparisons.

Altogether, our CAR-T cell functional atlas provides a resource to identify molecular determinants of efficacy, toxicity, and persistence in CAR-T therapy. Its open and expandable nature establishes a framework for continuous integration of new datasets and hypothesis generation that should contribute to accelerate the design of safer and more effective next-generation CAR-T therapies.

## DATA AVAILABILITY

All data in this study are publicly available. Statistics, resources and corresponding studies are listed in Supplementary Tables 1 and 2. The atlas pretrained model is broadly available to the community through scVI-hub at https://huggingface.co/sergiocamarap/Functional-cart-atlas-model, all processed data generated in this study have been deposited In Zenodo at https://doi.org/10.5281/zenodo.17213452 and the atlas is also available as an interactive Shiny application at https://wholebioinfo.shinyapps.io/shinyatlas/.

## CODE AVAILABILITY

All code used for data processing and visualization in this study is openly available at https://github.com/ML4BM-Lab/Functional-cart-atlas.

## ACKNOWLEDGEMENTS

This work was supported by the Spanish Ministry of Science, Innovation and Universities (MCIN/AEI, 10.13039/501100011033) through project PID2023-151980OB-I00, and PID2022-137914OB-I00, co-financed by the European Regional Development Fund (FEDER, “A way to make Europe”). Instituto de Salud Carlos III (ISCIII) through the Red de Terapias Avanzadas TERAV (RD21/0017/0009, RD21/0017/0019), RICORS TERAV Plus (RD24/0014/0010), and the Centro de Investigación Biomédica en Red de Cáncer (CIBERONC; CB16/12/00489, CB16/12/00369). The European Commission (T^2^EVOLVE, H2020-JTI-IMI2-2019-18, grant agreement No 945393 and EASYGEN, HORIZON-JU-IHI-2024-07, grant agreement No 101194710). Government of Navarra, Department of Industry (DIAMANTE, 0011-1411-2023-000074; 0011-1411-2023-000105) and Department of Health (GN2023/08; GN2024/04). “La Caixa” Foundation under the project code LCF/PR/HR24/52440011. Paula and Rodger Riney Foundation. Alberto Palatchi Foundation.

LJU. acknowledges support from a WIT grant (Marie Skłodowska-Curie Actions, Horizon 2020). ET acknowledges support from an AECC Clinico Junior Grant (CLJUN258694TAMA), and PRO from an AECC Clinico Senior Grant (CLSEN246328RODR), both funded by the Spanish Association Against Cancer (Asociacion Española Contra el Cancer, AECC). MH was supported by grant RYC2021-033127-I and NP was supported by grant RYC2021-032197-I, both funded by MICIU/AEI/10.13039/501100011033 and European Union NextGenerationEU/PRTR.; SCP from an FPU fellowship (FPU2023-00439); and from a CIMA AC fellowship.

We thank the authors of the original works that contributed to this atlas, with special gratitude to Zinaida Good, Kirk Gosik, Fernando Calero, Xiaonan Wang, Jos Melenhorst, Michael Green, and Zhiliang Bai for their invaluable support in gathering the necessary data.

## AUTHOR CONTRIBUTIONS

Conceptualization: SCP, PRM, JRRM, FP, MH. Formal analysis: SCP, NP, MECC. Data Curation: SCP, PRM, NP, LJU. Feedback: NP, LJU, GC, SRZ, PRO, ET, IO, NY. Visualization: SCP, PRM, JRRM, FP, MH. Writing: SCP, PRM, JRRM, FP, MH. Supervision: JRRM, FP, MH.

## COMPETING INTERESTS

The authors declare no competing interests.

## METHODS

### Acquisition of publicly available datasets

We considered publicly available studies analyzing CAR-T cells with single-cell sequencing as eligible for this project. In total, we acquired data from 13 studies^1–13^. When possible, raw sequencing data in FASTQ format was obtained to minimize batch effects. However, when it was unavailable, the count matrices provided by the authors were used (Supplementary Table 1). FASTQ files were processed into count matrices by aligning them against the human reference genome (GRCh38.p13) using the GENCODE release 42 annotation. For studies utilizing 10X Genomics technology, the 10X Cell Ranger suite (v7.1.0)^14^ was employed. In the case of studies using 3’scFTD-seq technology, the Drop-seq Core Computational tool (v2.5.1)^15,16^ was used, following the developers’ recommendations. Additionally, metadata including patient age, sex, therapy-related toxicities, response type, and other relevant clinical information was collected for all the samples included in the atlas (Supplementary Table 1).

### Identification of CAR^+^ T cells

This atlas was constructed exclusively from CAR^+^ T cells. These were distinguished from CAR^−^ T cells using the detection methods described by the original authors of each study (Supplementary Table 1 “CAR_Detection_Method”). For datasets in which the original CAR construct sequence was available, a custom reference incorporating the construct was generated, and cells with at least one detected CAR count were retained. Finally, to minimize potential noise in downstream analyses, samples with few CAR-expressing cells were excluded. Furthermore, the CAR gene was removed from all count matrices to avoid introducing bias into downstream analyses.

### Single-cell quality control

Count matrices were loaded into Seurat (v4.3.0.1)^17^ for quality control. Metrics such as number of detected genes, number of Unique Molecular Identifiers (UMIs), percentage of mitochondrial transcripts, and cellular complexity were manually inspected for each sample, then dataset-specific thresholds were applied. Cells lacking CD3 expression were excluded. Furthermore, artifacts commonly observed in droplet-based studies, such as doublets or empty droplets, were also considered, as their presence can bias biological interpretations and increase false discovery rates for rare populations. Doublets were identified using DoubletFinder (v2.0.3)^18,19^. Approximately 100 doublets consisting of T cells and erythrocytes (based on CD3 and of hemoglobin (e.g. HBA, HBB) gene expression) bypassed the threshold and were manually excluded. Empty droplets were detected using the *emptyDrops* function from DropletUtils (v1.14.2)^20^ for the 3’scFTD-seq datasets, consistent with the default method implemented in 10X Cell Ranger.

### Integration

Integration is a critical step when constructing an atlas from multiple sources, as it aims to mitigate technical biases while preserving the biological signal. To evaluate integration performance, six widely used tools were compared: scVI^21^, fastMNN^22^, Harmony^23^, Seurat RPCA^17^, STACAS^24^, and LIGER^25^. Benchmarking was performed using scGraph (v0.1.2)^26^, which quantifies the trade-off between batch correction and biological conservation (see Supplementary Note 1 for details). To ensure objectivity and minimize heterogeneity, this evaluation was carried out exclusively on healthy donor samples. Based on this comparison, scVI (v0.20.3) was selected. This Python method, based on deep neural networks, has been described as one of the top-performing tools for scRNA-seq, particularly in large and heterogeneous datasets with strong batch effects^27^ (Supplementary Note 1). Following the authors’ recommendations, raw counts were normalized by library size, log-transformed, and the analysis was limited to the 2,000 most variable genes, selected using the Seurat v3 flavor (span parameter = 0.6). Default scVI parameters were then applied for training and downstream processing. To enable the transition from the R to the Python environment, SeuratDisk (v0.0.0.9020) was used, and the subsequent analysis were conducted in Scanpy (v1.9.5)^28^. The final trained scVI model was exported into scVI-hub^29^.

### Annotation

Following integration, cells were meticulously annotated to ensure the reliability of downstream analyses. The integrated atlas was manually annotated using a set of curated marker genes: Cytotoxic/effector (*NKG7, GNLY, IFNG, PRF1, GZMK, GZMB*); activation (*CD69, CD27, CD28, HLA-DRA, TNFRSF9*); memory (*TCF7, CCR7, SELL*); regulatory T cells (*FOXP3, IL2RA, CTLA4*); proliferative (*ZWINT, MKI67, TUBA1B, TUBB*); monocyte-like T cells/IACs (*CD3*^*+*^, CD68, *CD14, LYZ*); and apoptotic cells (*CASP3, CASP8, BAD, FAS, BAK1, AKT1, CSE1L, TP53*). A group of cells enriched in ribosomal genes remained unannotated as it could not be confidently assigned to any of these categories.

### Validation of IACs as true cells

To rule out the possibility that monocyte-like T cells or ICANS-associated CAR-T cells (IACs) represented technical doublets between monocytes and T cells, artificial doublets were generated for comparison following the same strategy as the authors of the original report^3^. Specifically, two non-IAC cells were randomly selected, their raw count vectors summed up, and the result divided by two to approximate to the expression profile of a real doublet. This procedure was repeated iteratively to generate 6,000 artificial doublets. Artificial doublets were merged with the IACs population and processed with a similar pipeline, including normalization, log-transformation, selection of highly variable genes, dimensionality reduction (PCA), and UMAP visualization. The artificial doublets formed a single cluster distinct from the IACs cluster, supporting that IACs represent a genuine population rather than technical doublets.

### Differential abundance analysis with Milo

To investigate whether the composition of Infusion Product (IP) differed between clinical response groups differential abundance testing was applied using milopy (v0.1.1)^30^, the Python implementation of the Milo framework based on k-nearest neighbor (kNN) graphs. The analysis focused on IPs samples from complete responders (CR) and non-responders (NR) from patients aged over 40 years old, excluding *in vitro*-stimulated samples. A neighborhood graph was constructed from the scVI latent space (d = 30, k = 60), and neighborhoods were tested using the formula *~Age + Sex + Response*, with this last one as the variable of interest. Differential abundance was quantified as log fold change (logFC) between CR and NR, and statistical significance was assessed using SpatialFDR. Neighborhoods were annotated by their majority cell type label, note that those with <60% purity were excluded.

### Pseudo-bulk differential expression and pathway enrichment analysis

To characterize transcriptional changes, we applied dreamlet (v1.0.3)^31^ a pseudobulk framework based on precision-weighted linear mixed models, to aggregate single-cell expression profiles into pseudo-bulk matrices stratified by relevant biological conditions. Single-cell counts were aggregated by sample (sample_id = Product_norm), while the biological stratum (cluster_id) was adapted to each analysis: time points in CR CD8^+^ cytotoxic T cells (Infusion Product, <2 weeks, 2–3 months, >3 months), clinical response comparison in CD8^+^ cytotoxic T cells (<2 weeks vs. IP), IACs (IP vs. post-infusion; and IACs vs. the rest CAR-T cells), and CAR type group (BCMA vs. CD19). Gene set enrichment analyses were performed with zenith (v1.4.2)^32^ using some Enrichr-sourced collections (GO Biological Process 2023, KEGG 2021 Human, Reactome 2022, and WikiPathways 2023 Human) together with manually curated CAR-T signatures (activation, tonic signaling, cytotoxicity, memory, exhaustion, and metabolic programs). For selected contrasts (e.g., IACs post-infusion vs. IP), complementary enrichment dot plots were generated with clusterProfiler (v4.2.2)^33^ to visualize GO Biological Process enrichments. To account for multiple hypothesis testing, p-values were adjusted using the Benjamini-Hochberg procedure, and results are reported as FDR-adjusted values.

### AUCell scoring of IL-10 signature

To evaluate the activity of IL-10, we applied the AUCell package (v1.30.1)^34^ to quantify enrichment of a curated IL-10 gene signature (Table S4). AUCell estimates the relative activity of a gene set in single cells by calculating the are under the curve (AUC) of the ranked gene expression distribution. We restricted the analysis to IP cells from patients with available clinical response annotations (CR vs. NR). Finally, AUC scores were computed for individual cells across cell types and contrasted between patients with different clinical responses.

### Differential proportion analysis of IP

Differences in the relative abundance of specific cell subsets were assessed using scProportionTest (v0.1.2)^35^. This framework compares cell type frequencies between groups by permutating sample labels to generate a null distribution and applying bootstrapping to estimate confidence intervals for the observed effect sizes. We applied this approach to investigate demographic and clinical influences on IP composition, including sex (female vs. male), age (40-60 vs. >60 years old), and clinical response groups (CR vs. NR). Reported p-values correspond to permutation-derived significance estimates with 10,000 permutations and 10,000 bootstraps.

### Integration of new datasets (scArches-scANVI)

New datasets were integrated using scArches (v0.6.1) in combination with scANVI^36^, following the pipeline recommended by the authors. Specifically, the reference atlas was trained with 2,000 highly variable genes using default hyperparameters. A scANVI model was then initialized from the trained scVI reference and trained for 20 epochs under default settings. Query datasets were subsequently integrated through the transfer learning framework implemented in scArches, with dropout frozen and training for 100 epochs with a batch size of 256.

### Interactive atlas visualization

An interactive web application was generated using ShinyCell (v2.1.0)^37^ to enable researchers without advanced bioinformatics expertise to explore the atlas. The application was generated from the final AnnData object and includes precomputed UMAP embeddings, curated cell type annotations, and associated metadata (Supplementary Table 1). It allows users to interactively explore the atlas, such as visualizing gene expression or cell type distributions across samples. The link to the interactive web is https://wholebioinfo.shinyapps.io/shinyatlas/.

## Notes

https://huggingface.co/sergiocamarap/Functional-cart-atlas-model

https://doi.org/10.5281/zenodo.17213452

https://wholebioinfo.shinyapps.io/shinyatlas/

https://github.com/ML4BM-Lab/Functional-cart-atlas

### Competing Interest Statement

The authors have declared no competing interest.

