## Extended data for "An open CAR-T single-cell atlas to enable in-depth characterization and rational engineering of CAR-T products"

### <u>SUPPLEMENTARY MATERIAL</u>

**SUPPLEMENTARY TABLES TITLE…………………………………………………………………………………………..……2**

**SUPPLEMENTARY NOTE…………………………………………………………………………………………………..….3**

**EXTENDED DATA FIGURES……………………………………………………………………………………………….….4**

**SUPPLEMENTARY TABLES TITLE**

**Table S1. CAR-T cell atlas complete metadata.**

**Table S2. Metadata summary across individual studies.**

**Table S3. scGraph metrics scores for each integration method.**

**Table S4. IL-10 signaling pathway associated genes.**

**Table S5. Wilcoxon test results comparing IL-10 signature AUC scores between CR and NR across CAR-T cell subsets.**

**Table S6. Fisher's exact test of the association between IL-10 expression and clinical response (CR vs. NR) at cell level.**

**Table S7. Dreamlet results for pathway differences in $CD8^+$ cytotoxic cells at early time points between CR and NR/PR.**

**Table S8. Wald test results for pathway differences in $CD8^+$ cytotoxic cells at early time points between CR and NR/PR.**

**Table S9. Differential expression of inflammatory genes in IACs between IP and post-infusion samples (Mann–Whitney U test).**

**Table S10. Contingency table of IP IACs vs ICANS severity at the patient level, with associated Fisher's exact test statistics.**

**Table S11. Metadata from the newly integrated dataset.**

**Table S12. Wald test results for pathway differences between BCMA- and CD19-targeted CAR-T cells at IP and mid-time after infusion.**

**SUPPLEMENTARY NOTE**

**Sup. Note 1 – Benchmarking of dataset integration methods**

Integration of heterogeneous scRNA-seq datasets is a known critical step in the construction of atlases. Inappropriate integration may lead either to the removal of relevant biological signal or, in the other hand, to the persistence of technical artifacts that may obscure the underlying biology. However, no gold-standard method exists that can handle all integration scenarios effectively, as the results from these tools depend heavily on the type of data used and the project's objectives. Given the importance of this step for downstream analysis, we evaluated six of the most used integration methods (LIGER, scVI, STACAS, Harmony, fastMNN, and Seurat RPCA). Integration performance was assessed with scGraph[1], which quantifies the trade-off between batch correction and biological conservation.

As expected, the merged dataset, showed the lowest scores across all metrics, reflecting strong batch-driven structure (Corr-Weighted = 0.08). Among integration methods, LIGER achieved the highest scores (Corr-Weighted = 0.66), leaving scVI as the second-best method overall (Corr-Weighted = 0.57) (Fig. S2, Table S3).

Although LIGER performed slightly better on the scGraph metrics, we ultimately selected scVI as the integration framework for our atlas. This decision was made based on the evidence that an external benchmarking shows. In particular, the scIB[2] benchmark reported that scVI consistently ranks among the best-performing methods in scenarios involving large number of cells and complex batch structures. In these challenging settings, scVI was shown to effectively reduce batch effects while retaining biologically meaningful variation, performing particularly well in immune cell datasets, that are the closest to the characteristics of our study. Moreover, it is said that compared with alternative approaches that either over-correct and tend to remove biological signal (e.g., LIGER, Seurat) or fail to achieve sufficient batch mixing, scVI provides a more balances integration. The final recommendation from the authors is to use scVI in the absence of labels, especially if you are integrating large datasets.

Together, these independent findings, reinforced our empirical results and validate scVI as the most suitable method for integrating the heterogeneous datasets that are included in this study.

**EXTENDED DATA FIGURES**

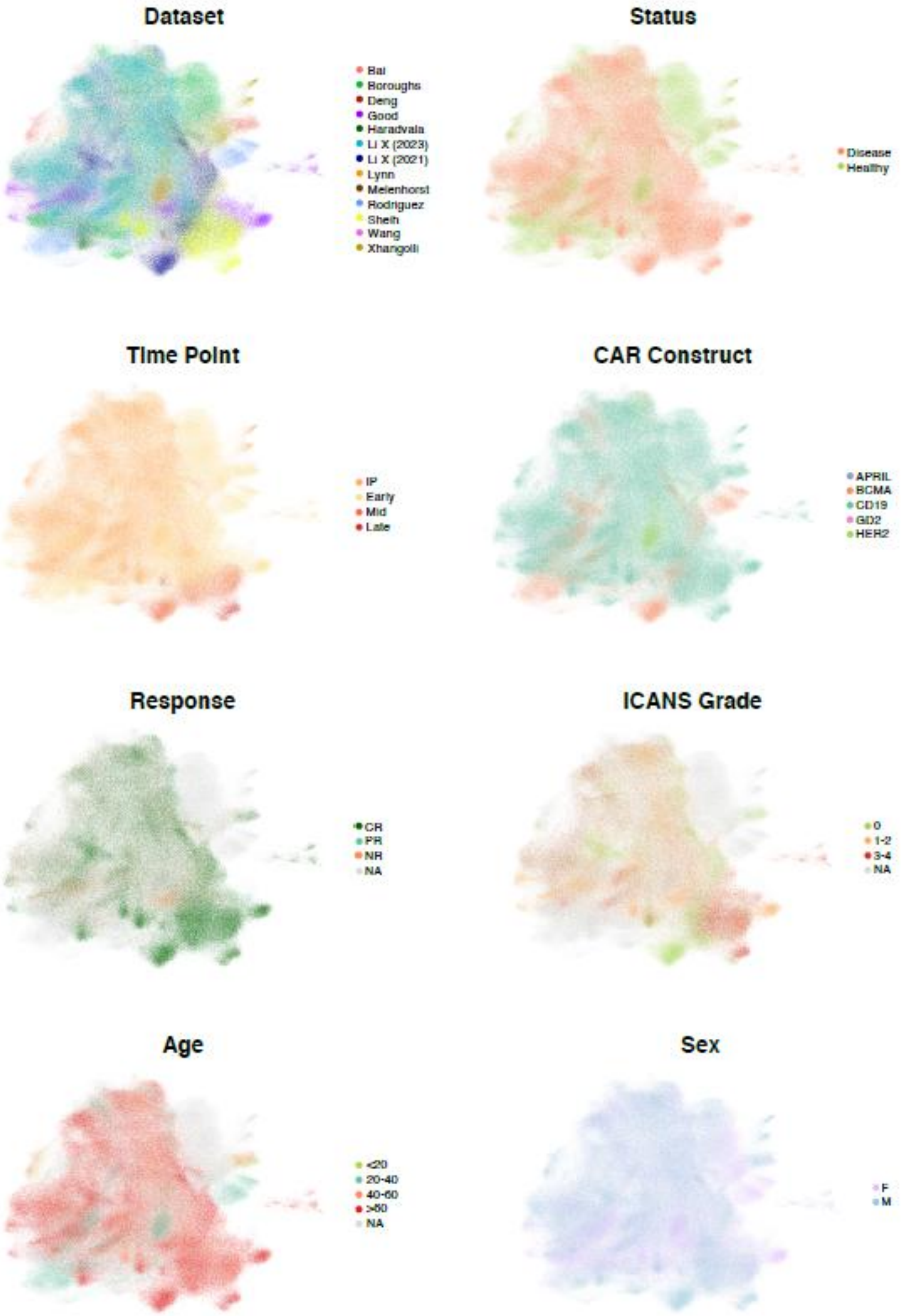


**Extended Data Fig. 1. CAR-T cell atlas UMAP colored by key clinical and experimental features.** CAR-T cells are colored according to dataset of origin, status (disease, healthy), collection time point (infusion product [IP], early [<2 weeks], mid [2 weeks–3 months], late [>3 months]), CAR construct (APRIL, BCMA, CD19, GD2, HER2), clinical response (CR, PR, and NR), ICANS grade (0, 1–2, and 3–4), patient's age (<20, 20–40, 40–60, and >60), and sex (female [F], male [M]). NA: Not applicable.

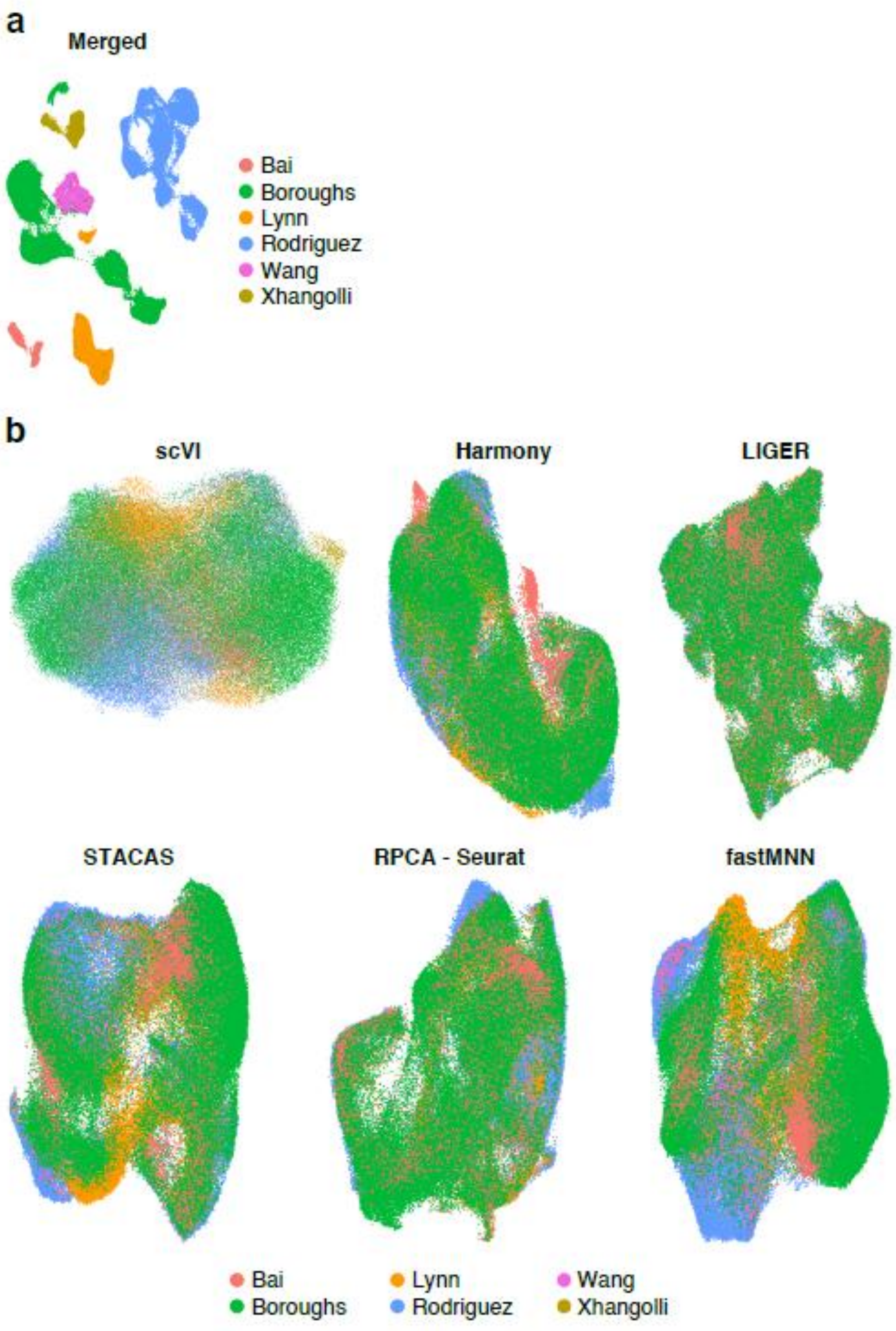


**Extended Data Fig. 2. Evaluation of CAR-T cell atlas dataset integration methods. a,** UMAP of the CAR-T cell atlas merged datasets without integration, showing that integration was needed. **b,** Six integration approaches were evaluated (scVI, Harmony, LIGER, STACAS, Seurat RPCA, and fastMNN), and scVI was finally selected. UMAP showing the results of integration of all samples for each method are depicted.

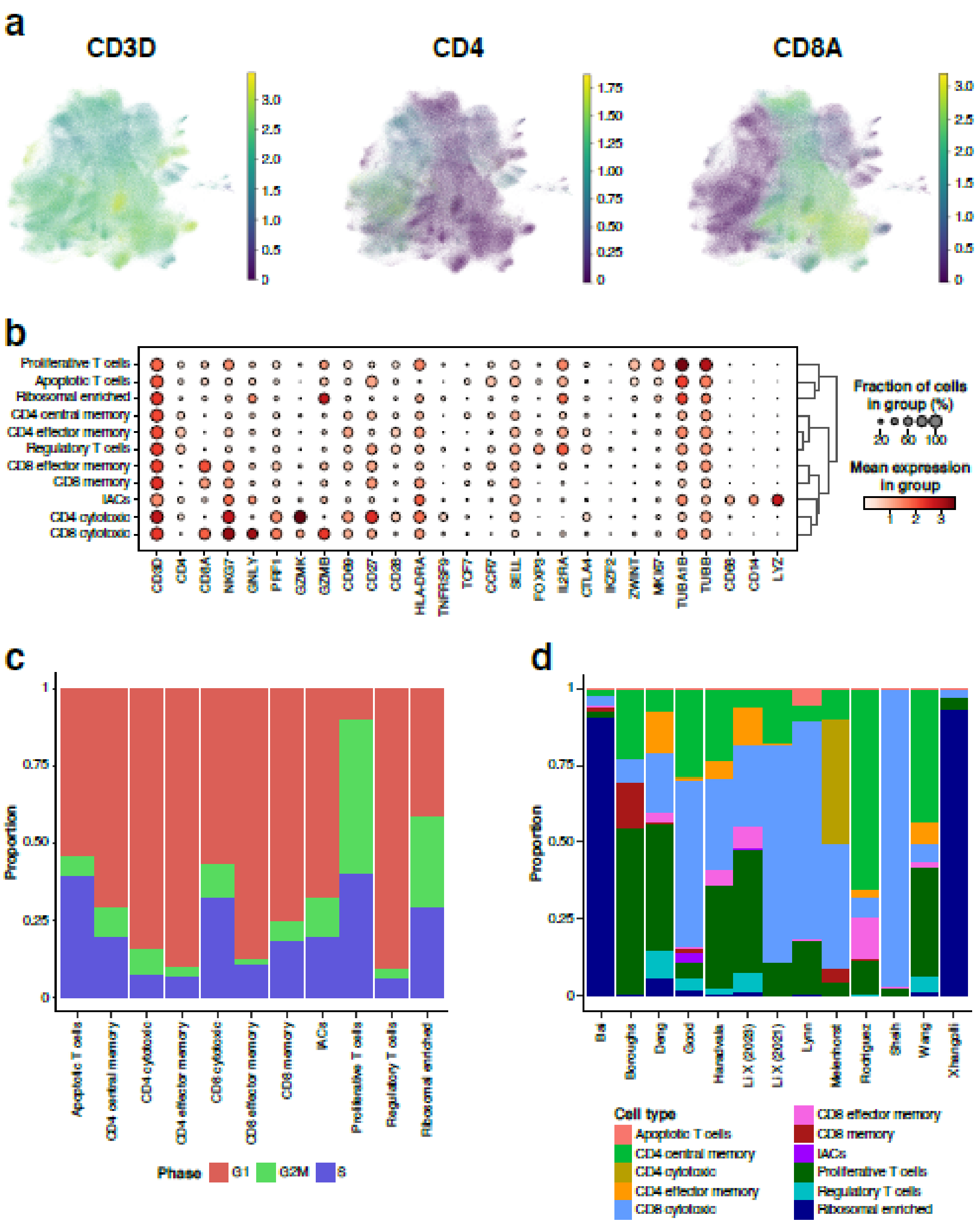


**Extended Data Fig. 3. Annotation of T cell populations in the integrated CAR-T cell atlas. a,** UMAP colored by T cell main genes (CD3D, CD4, and CD8A) expression. **b,** Dot plot of the expression of most relevant marker genes used for annotation of T cell populations. **c,** Distribution of cell cycle phases (G1, S, and G2/M) across annotated T cell populations. **d,** Distribution of each annotated T cell across the different studies included in the atlas.

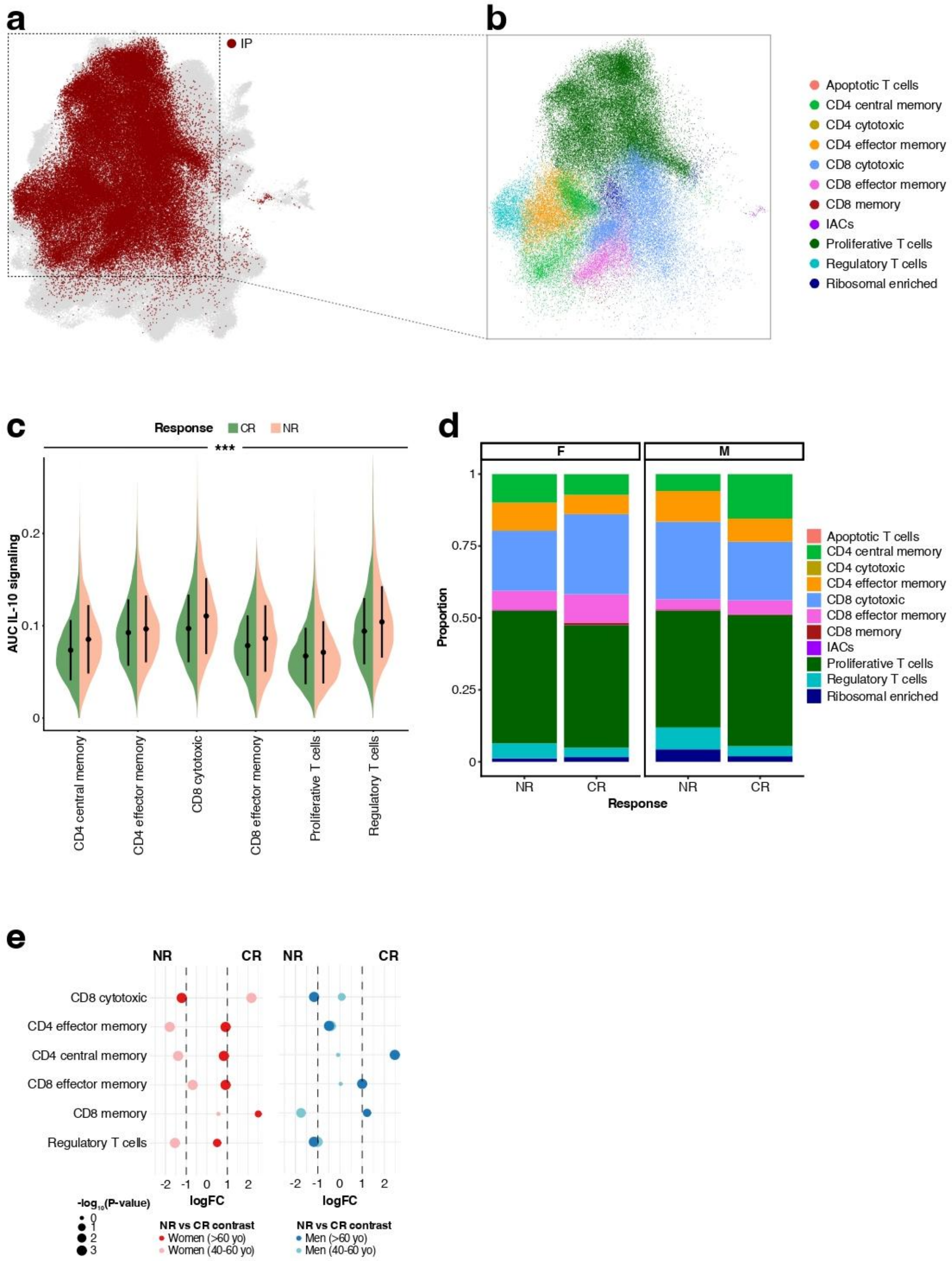


**Extended Data Fig. 4. Transcriptomic characterization of infusion products. a,** A subset of CAR-T cells from IP (red) was selected for differential abundance testing (CR and NR patients >40 years old). **b,** IP cells subset colored by T cell population. **c,** Comparison of AUC scores of IL-10 signaling signature within CR and NR patients across the different cell compartments. **d,** Phenotype proportions in NR and CR patients divided by sex (female [F] and male [M]). **e,** Permutation test for NR and CR patients divided by sex and age (40-60 yo, >60 yo). Wilcoxon test was used for c. ***P < 0.001.

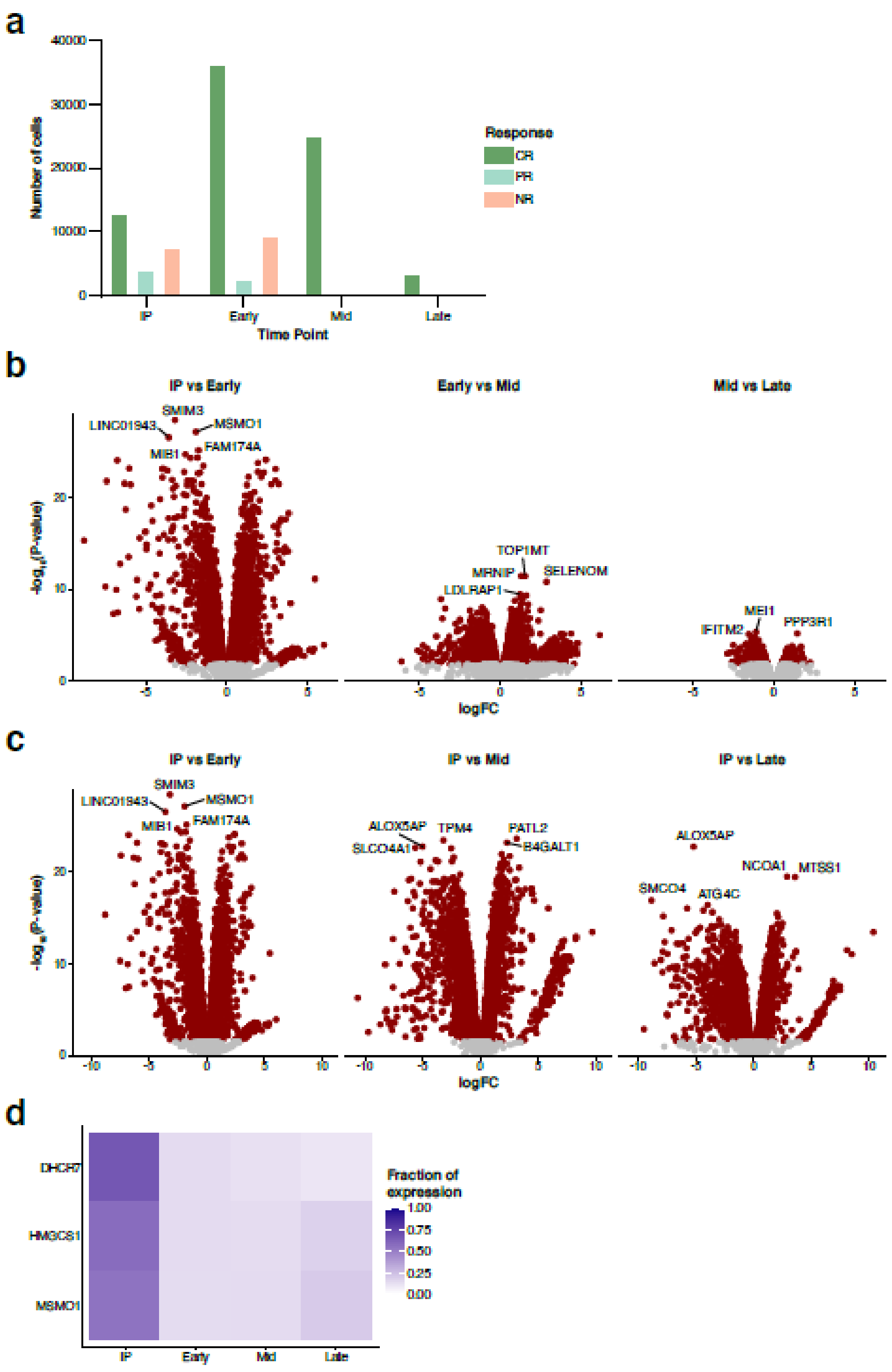


**Extended Data Fig. 5. Molecular changes of CAR-T cells in CR patients after infusion.** We focused on the transcriptional changes in $CD8^+$ cytotoxic CAR-T cells comparing IP with post-infusion time points. **a,** Number of $CD8^+$ cytotoxic cells across time points divided by clinical response (CR, PR, NR). **b,** Volcano plots of differentially expressed genes between consecutive time points (IP vs early; early vs mid; mid vs late). **c,** Volcano plots of differentially expressed genes comparing each time point against IP. **d,** Cholesterol biosynthesis genes (*DHCR7*, *HMGCS1*, and *MSMO1*) showed upregulation in IP compared to post-infusion time points.

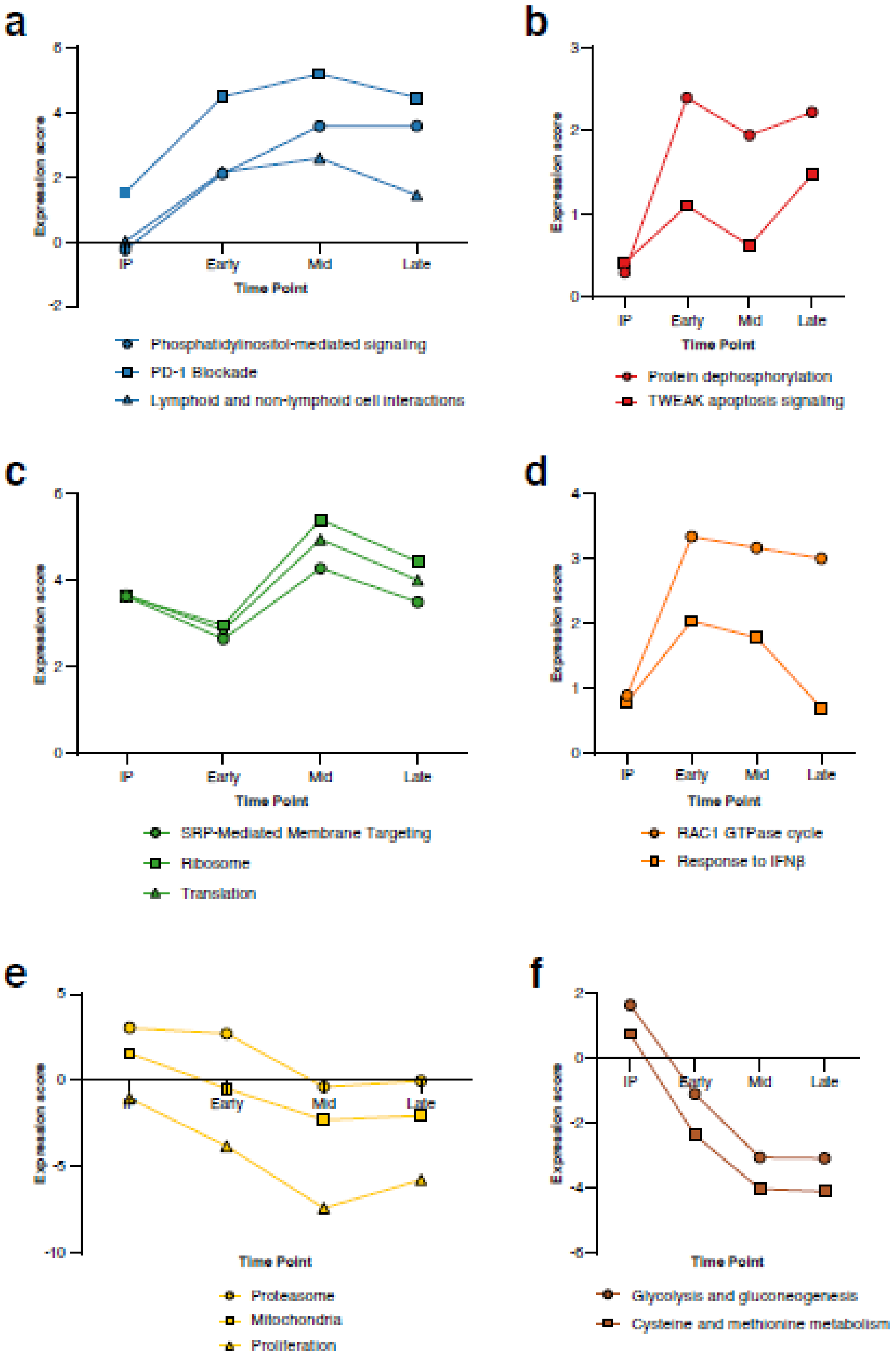


**Extended Data Fig. 6. Temporal dynamics of pathway signatures in $CD8^+$ cytotoxic CAR-T cells from CR patients.** Pathways are grouped into subpanels according to their trajectories. **a, b,** Pathways upregulation in early-mid points and maintenance at later time points. **c,** Downregulation in short time points followed by increase of expression at later stages. **d,** Upregulation in short-mid time points followed by recovery at late timepoint. **e, f,** Continuous decrease along time.

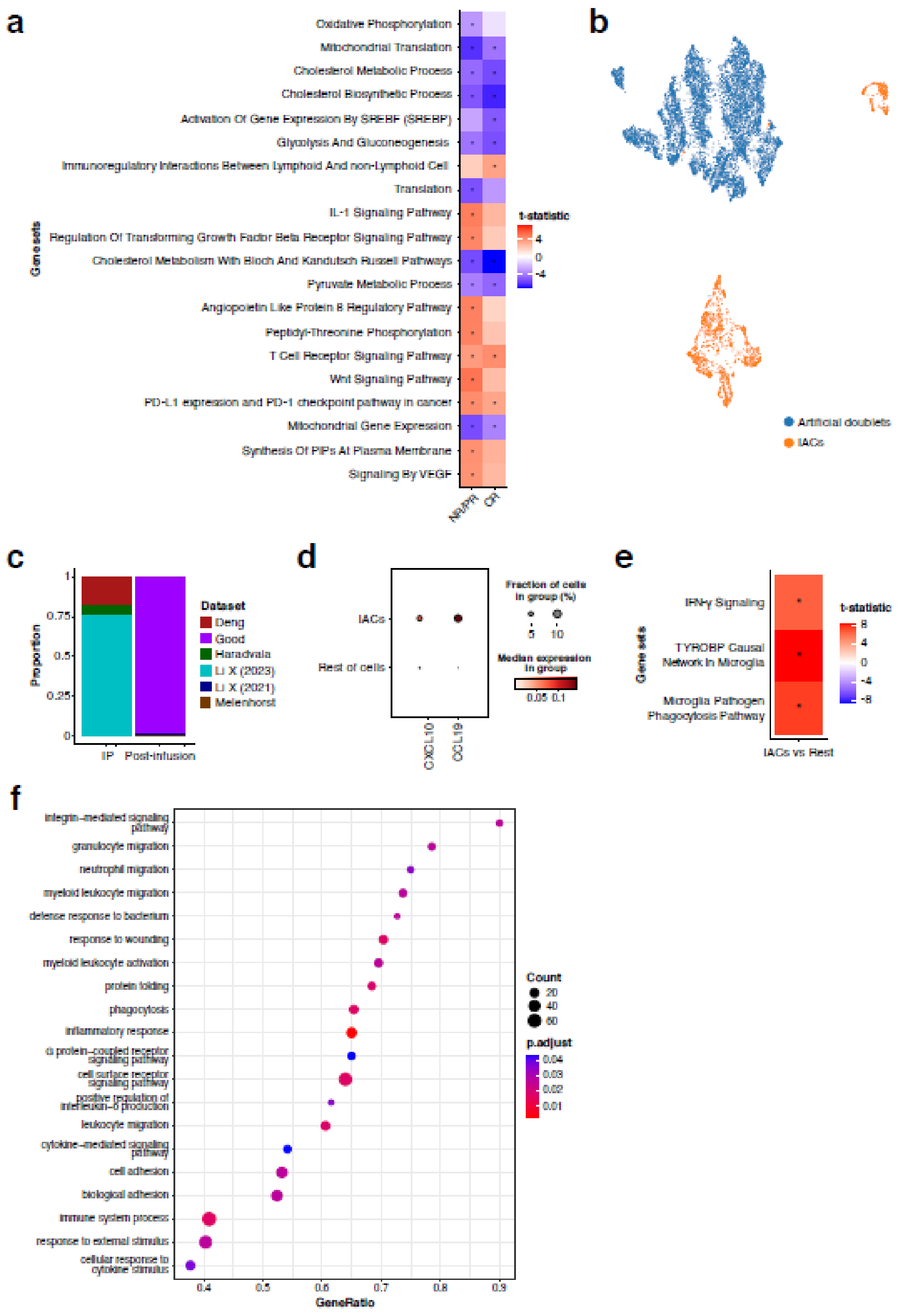


**Extended Data Fig. 7. Response-associated pathways in CD8+ cytotoxic cells and characterization of ICANS-associated CAR-T cells (IACs). a,** Pathways differentially expressed between CD8$^{+}$ cytotoxic cells from NR/PR compared to CR at early time point. **b,** Artificial doublets were generated to compare with IACs. UMAP showed that they cluster separately, confirming that IACs are not doublets. **c,** Proportion of IACs across datasets, stratified by IP and post-infusion samples. **d,** Expression of inflammatory genes *CXCL10* and *CCL19* in IACs compared with the rest of cells in the atlas. **e,** Enriched pathways in IACs compared with the rest of cells in the atlas. **f,** GSEA between IACs from IP and post-infusion.

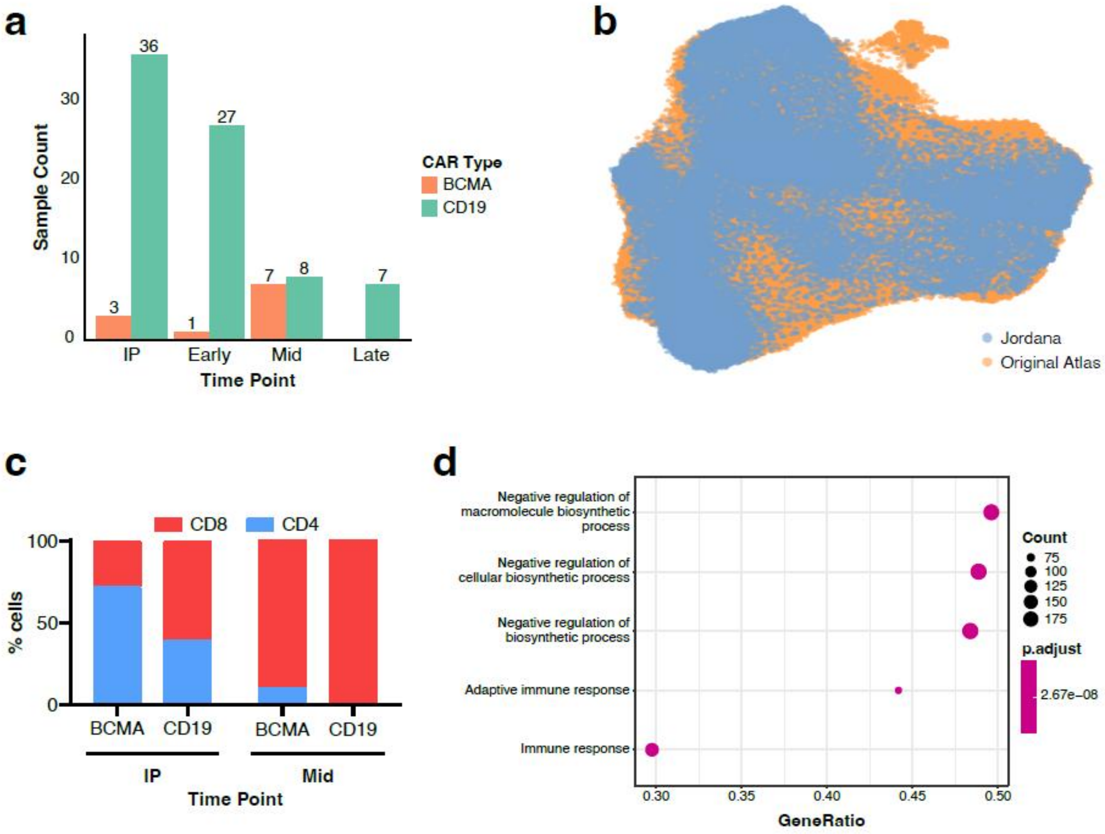


**Extended Data Fig. 8. Integration of a new in-house BCMA dataset and comparison with CD19 CAR-T cells. a,** Distribution over time of BCMA and CD19-targeting CAR-T samples after integration. **b,** UMAP with the new dataset (Jordana) integrated with the original atlas. **c,** Proportion of $CD4^{+}$ and $CD8^{+}$ CAR-T cells in BCMA and CD19 samples at IP and mid time point. **d,** GSEA of pathway enrichment differences between BCMA- and CD19-targeting CAR-T cells.
